# Interactions of anti-COVID-19 drug candidates with multispecific ABC and OATP drug transporters

**DOI:** 10.1101/2020.11.21.392555

**Authors:** Ágnes Telbisz, Csilla Ambrus, Orsolya Mózner, Edit Szabó, György Várady, Éva Bakos, Balázs Sarkadi, Csilla Özvegy-Laczka

## Abstract

In the COVID-19 epidemic, several repurposed drugs have been proposed to alleviate the major health effects of the disease. These drugs are often applied together with analgesics or non-steroid anti-inflammatory compounds, and co-morbid patients may also be treated with anticancer, cholesterol-lowering or antidiabetic agents. Since drug ADME-tox properties may be significantly affected by multispecific transporters, here we examined the interactions of the repurposed drugs with the key human multidrug transporters, present in the major tissue barriers and strongly affecting pharmacokinetics. Our in vitro studies, using a variety of model systems, explored the interactions of the antimalarial agents chloroquine and hydroxychloroquine, the antihelmintic ivermectin, and the proposed antiviral compounds, ritonavir, lopinavir, favipiravir and remdesivir with the ABCB1/Pgp, ABCG2/BCRP and ABCC1/MRP1 exporters, as well as the OATP2B1 and OATP1A2 uptake transporters. The results presented here show numerous pharmacologically relevant transporter interactions and may provide a warning for the potential toxicities of these repurposed drugs, especially in drug combinations at the clinic.

## 1. Introduction

During the outbreak of the COVID-19 pandemic, based on *in vitro* experimental studies, a number of potential antiviral drugs have been proposed to reach the clinic. These potential treatments were rapidly brought to the attention of the medical community and the general public by the media, while in many cases drug evaluation agencies could not properly investigate the pharmacokinetics, the potential risks and benefits. Still, clinicians and thousands or even millions of lay people started a compassionate use of the advocated off-label compounds.

The most notorious example is the wide range off-label use of the antimalarial agents, **chloroquine and hydroxychloroquine**, in some cases together with zinc or the wide-spectrum antibacterial agent, azithromycin. Chloroquine (CQ) and the less toxic analog hydroxychloroquine (HCQ) are efficient antimalarial drugs, increasing the endosomal pH both in the parasites and the host cells. HCQ is also clinically used to treat autoimmune diseases, such as systemic lupus erythematosus (SLE) and rheumatoid arthritis[1]. Both of these compounds were found to have major toxicities, especially by having additive effects with other drugs prolonging the cardiac QT interval or causing hypoglycemia. CQ and HCQ were both reported to be moderate inhibitors of CYP2D6 and the ABCB1/Pgp transporter. Interestingly azithromycin has a similar toxicity as CQ and HCQ, that is an additive effect with other drugs that prolong the QT interval[2].

The potential use of CQ and HCQ in COVID-19 was initiated by *in vitro* studies, in which both CQ and HCQ were found to inhibit the fusion of SARS-CoV-2 and the host cell membranes, inhibit the glycosylation of the cellular ACE2 receptor (the binding site of SARS-CoV-2), and block the transport of SARS-CoV-2 from early endosomes to endolysosomes[3–5]. However, by now CQ and HCQ, with or without azithromycin, have been studied in multiple clinical trials for the treatment of COVID-19, and the use of these agents has not been approved by the EMA or FDA. A recent statement of the NIH COVID-19 Treatment Guidelines Panel strongly advises against the use of CQ or HCQ because of ineffectivity, and the findings that a concomitant use of HCQ and azithromycin can prolong the QT interval, and associates with increased odds of cardiac arrest[6,7].

Another proposed anti-COVID-19 agent, **ivermectin**, is a member of the class of avermectins which are highly active broad-spectrum, anti-parasitic agents and are used to treat several neglected tropical diseases, including onchocerciasis, helminthiases, and scabies. The target of the antiparasitic action of ivermectin is a glutamate-gated chloride channel and a GABA receptor, specific for some invertebrates. However, in mammals, ivermectin may also inhibit GABAergic neurotransmission by promoting the release of GABA and acting as a GABA receptor agonist. This potentially neurotoxic drug is absorbed in humans only in a very small fraction, and this low level ivermectin absorption is mainly caused by an active extrusion in the intestine by the ABCB1/Pgp transporter. ABCB1, and probably other ABC transporters (especially ABCG2) in the blood-brain barrier (BBB) also have a significant role in protecting the mammalian CNS against toxic ivermectin penetration. In mouse Pgp-knock-out models, in the natural Pgp-knock-out Collie dogs, and also in some humans with low level ABCB1/Pgp expression, ivermectin exerts major neurotoxicity[8,9]. In addition, both ABCB1 and several other multispecific transporters have been shown to be inhibited by micromolar concentrations of ivermectin[10–12], thus ivermectin may influence the pharmacokinetics of several drugs or toxic compounds.

Ivermectin was suggested to have anti-CoV-2 effects based on *in vitro* studies, as this compound inhibits the cellular importin alpha/beta-1 nuclear transport proteins, thus in cell cultures reduces the replication of a wide range of viruses[13–15], including SARS-CoV-2[16,17]. However, pharmacokinetic and pharmacodynamic studies suggest that achieving the ivermectin plasma concentrations necessary for the *in vitro* antiviral efficacy (about 2-10 μM), is highly toxic and would require administration of oral doses up to 100-fold higher than those approved for use in humans[18]. Therefore, in spite of some anecdotal clinical results, ivermectin is not approved for the treatment of any viral infection, including SARS-CoV-2 infection according to the NIH COVID-19 Treatment Guidelines (NIH-CTGP). Moreover, the FDA issued a warning on April 2020 that ivermectin should not be used to treat COVID-19 in humans.

Several approved and clinically effective HIV protease inhibitors, including **ritonavir and lopinavir**, based on their potential for inhibiting other viral proteases, were also entered into clinical trials as anti-COVID-19 agents. The replication of SARS-CoV-2 depends on the cleavage of viral polyproteins into an RNA-dependent RNA polymerase and a helicase, and two proteases, 3CLpro and PLpro are responsible for this cleavage. *In vitro*, lopinavir and ritonavir were found to inhibit 3CLpro, and thus reduce SARS-CoV-2 replication[19]. Still, these drugs have a poor selectivity index, indicating that higher than tolerable levels of the drug might be required to achieve meaningful inhibition *in vivo*. Therefore, the current NIH-CTGP recommends against using lopinavir/ritonavir or other HIV protease inhibitors for the treatment of COVID-19, except in a clinical trial. The main cause of this negative recommendation is based on the pharmacodynamics of lopinavir/ritonavir, showing that these drugs are unlikely to reach *in vivo* concentrations that can inhibit the SARS-CoV-2 proteases[20]. Also, lopinavir/ritonavir did not show efficacy in a moderately sized randomized controlled trial in patients with COVID-19. The NIH-CTGP warns that lopinavir/ritonavir is a potent inhibitor of cytochrome CYP3A, thus co-treatment with numerous drugs metabolized by CYP3A may result in increased toxicities. Ritonavir induces CYP1A2 and inhibits CYP3A4 and CYP2D6, thus may cause serious drug-drug interactions. Ritonavir and lopinavir are inhibitors of ABC multidrug transporters[21,22] both *in vitro* and *in vivo*[23], and based on studies with labeled ritonavir and lopinavir, these agents are transported substrates of ABCB1/Pgp[24,25], while not of ABCG2/BCRP[26,27].

Recently developed antiviral agents, although found to be minimally effective in earlier clinical studies and not approved for general use, have also been introduced into COVID-19 clinical trials. **Favipiravir**(Avigan) is a pyrazinecarboxamide derivative antiviral agent, a prodrug that is metabolized to its active form, favipiravir-ribofuranosyl-5'-triphosphate (favipiravir-RTP) and can be used both in oral and intravenous formulations. The mechanism of its actions is thought to be a selective inhibition of viral RNA-dependent RNA polymerase and functions as a chain terminator of the viral RNA, thus reduces the viral load. Favipiravir was approved in Japan against special cases of influenza, although it has not been effective in primary human airway cells, questioning its efficacy[28].

Favipiravir degradation is not based on the cytochrome P450 system but occurs by aldehyde oxidase (AO) and xanthine oxidase. Therefore, a co-administration of the drug with AO inhibitors such as cimetidine, tamoxifen, phenothiazines, verapamil, amlodipine, nifedipine, loratidine, cyclobenzaprine, ondansetron, ketoconazole, could increase the level of favipiravir and its metabolites. Since favipiravir was found to inhibit CYP2C8, co-administration with CYP2C8 substrates e.g. paclitaxel, repaglinide and rosiglitazone may pose a risk for increased drug effects. Drugs which have been also indicated to interact with favipiravir are chloroquine, diclofenac, digoxin, doxorubicin, edoxaban, fluorouracil, ibuprofen, hydrocortisone, and hydroxychloroquine. Since favipiravir is likely to be co-prescribed with acetaminophen (paracetamol), it is to note that favipiravir inhibits acetaminophen sulfate formation. Faviparivir has been shown to have a teratogenic effect in animals[29]. Currently the use of favipiravir in treating COVID-19, although with only limited clinical data for efficiency in this disease, has been approved in China, India, and for emergency use in Japan, while this agent remains unapproved in Europe and the USA.

**Remdesivir**(Veklury) is a broad-spectrum antiviral medication for intravenous injection provided in a sulfobutylether-β-cyclodextrin (SBECD) complex. Remdesivir (RDV) itself is a prodrug, which after entering the cells is converted to a mono-phosphate by esterases and a phosphoamidase, then further phosphorylated to an active metabolite triphosphate by nucleoside-phosphate kinases. This latter compound is a ribonucleotide analogue inhibitor of viral RNA polymerase. Remdesivir was originally developed to treat hepatitis C and was also tested against Ebola virus disease, but found to be ineffective for these viral infections. However, based on initial clinical trials, remdesivir has been authorized for emergency use in COVID-19 in the US, India and Singapore, and approved for use in Japan, the European Union and Australia for patients with severe symptoms. In October 2020, FDA authorized the clinical use of remdesivir against COVID-19, while the WHO still does not support its use in this disease. Preclinical studies indicated that remdesivir is not a substrate of CYP2C8, CYP2D6, or CYP3A4, and is not “significantly” transported by ABCB1/Pgp or the OATP type drug transporters[30], while detailed studies for remdesivir-drug interactions are not available as yet. Remdesivir is apparently an *in vitro* substrate of CYP2C8, 2D6, and 3A4, while *in vivo* remdesivir metabolism is suggested to be dominated by hydrolase activity. Remdesivir at the clinic is applied in the form of a sulfobutyl ether beta-cyclodextrin (SBECD) complex thus, in addition to remdesivir, we have also used this cyclodextrin complex in our *in vitro* studies.

In order to explore the multispecific drug transporter interactions of the above repurposed drugs, here we performed a detailed *in vitro* study on their potential interaction with some key multidrug transporters. We focused on the potential role of the transporter-drug interactions in the important tissue barriers, especially the intestinal epithelium and the blood-brain barrier (BBB) endothelial cells. (see http://www.fda.gov/Drugs/DevelopmentApprovalProcess/DevelopmentResources/DrugInteractionsLabeling/ucm093664.htm#major (2017)).

The ABCB1/Pgp and the ABCG2/BCRP efflux transporters are key proteins involved in the xenobiotics extrusion in the intestine and in the BBB, while they are also involved in toxin and drug metabolism in the liver and kidney[31–33]. In addition to some endogenous substrates, including conjugated metabolites, steroids or uric acid, there is a wide range of environmental and food-related toxic molecules actively transported by these proteins, thus they are major players in drug-drug interactions. While in most model animals Pgp is the major drug transporter in the BBB, in primates and humans ABCG2 seems to be the key protective transporter in this tissue[34–36]. In addition, ABCC1 may have an important role in the blood-cerebrospinal barrier to avoid CNS toxicity[37].

Organic anion-transporting polypeptides (OATPs) are members of the Solute Carrier (SLC) family and some of them (e.g. OATP1A2, 1B1, 1B3 and 2B1) are well documented multispecific plasma membrane xenobiotic and drug transporters that mediate the cellular uptake of a large variety of organic molecules (“uptake transporters”)[38]. OATP1B1 and OATP1B3 have a key role in the liver drug metabolism, while OATP1A2 and OATP2B1 are widely expressed in human tissues[39]. These latter transporters are highly expressed in the intestine, while OATP1A2 is the key drug uptake transporter expressed in the luminal side of BBB endothelial cells, promoting drug uptake from the blood into the CNS[40–42].

Therefore, OATPs 1A2 and 2B1 are important determinants of the pharmacokinetics of numerous drugs, either being substrates or inhibitors of these transporters. Based on both *in vitro* and clinical data, OATP1A2 and OATP2B1 are key players in the intestinal uptake of a large variety of clinically important drugs, including statins, fexofenadine, sulfasalazine, steroids or telmisartan, while may be significantly inhibited by chemotherapeutics or antiviral compounds[42]. In addition, they contribute to the tissue penetration of numerous endogenous substrates, including steroid and thyroid hormones, prostaglandins, bile acids, or bilirubin[43].

Based on their important role in tissue barriers and drug pharmacokinetics, we have studied the potential inhibition of the function of ABCB1, ABCG2, ABCC1, OATP2B1 and OATP1A2, by the clinically applied anti-COVID-19 agents. In these experiments we have used an array of *in vitro* functional transporter assays which together may provide important new information regarding the potential ADME-tox properties of these agents.

## 2 Materials and Methods

*Materials:* All basic laboratory reagents were from Sigma Aldrich. The 5D3 antibody was a kind gift of B. Sorrentino, St. Jude Children Hospital, Memphis, USA, remdesivir and Remdesivir-SBECD were kind gifts of Lajos Szente, Cyclolab Ltd, Budapest, Hungary. Vesicular assay membranes and components were obtained from SOLVO Biotechnology, Hungary (https://www.solvobiotech.com/services/categories/)

For the *ABC transporter assays* we have used both cell-based and membrane based functional assays.

*For assaying the function of the ABC transporters in cell-based assays* we applied the stable cell lines expressing these transporters: ABCG2 PLB-985, ABCB1 PLB-985, ABCC1 HL-60, and their parental lines that lack significant ABC transporter expression[44]. The respective transport activities were assayed by measuring the cellular fluorescence of the respective transported substrates Phengreen (ABCG2) or Calcein (ABCB1 and ABCC1)[44]. For the ABCG2 transport assay, we have also used HeLa cells stably expressing the wild-type or Q141K polymorphic variant of the ABCG2 protein (see below).

*For studying ABCB1 and ABCC1 transport*, 5×10^4^ parental control or transporter containing cells were incubated with 0.25μM Calcein-AM (C3100MP ThermoFisher Scientific, Walthman, MA, USA) in phosphate buffered saline (PBS) containing 1g/L D-glucose (DPBS) for 40 minutes at 37°C. The test compounds were applied in 0.2-50 μM, as specific transporter inhibitors 0.25 μM tariquidar (TQ, was a kind gift from Dr. S. Bates (NCI, NIH)) for ABCB1, and 10 μM indomethacin (IM, I0200000, Sigma-Aldrich-Merck, St. Louis, USA) were used. The cells were kept on ice until the flow cytometry measurements. All experiments were performed in triplicates and in at least three biological parallels.

*For examining the function of ABCG2*, 5×10^4^ parental control or ABCG2 transporter expressing cells were incubated with 0.25 μM PhenGreen-SK diacetate (P14313, ThermoFisher Scientific, Walthman, MA, USA) in DPBS containing 1μM EDTA, for 40 minutes at 37° C. The test compounds were applied in 0.2-50 μM, as specific transporter inhibitor 2.5 μM Ko143 (3241, Tocris Bioscience, Bristol, UK) was applied. The cells were kept on ice until the flow cytometry measurements. All experiments were performed in triplicates and in at least three biological parallels.

Cellular fluorescence reflecting transport activity was measured by Attune Nxt cytometer (Thermo Fischer Scientific, Waltham, MA, US) equipped with a blue (488 nm) laser. The PhenGreen (PG) or the Calcein signal was detected in the BL1 channel (emission filter: 530/30 nm). Data analysis was performed using the Attune Nxt Cytometer Software v3.1.2 (Thermo Fischer Scientific, Waltham, MA, US). Inhibition of transporter activity was calculated based on the fluorescence signal in the presence of the test compound, relative to that seen with the specific inhibitors providing maximum inhibition. Higher concentrations of some of the investigated compounds altered the fluorescence of Calcein also in the parental cells by an ABC transporter independent way (parental cells have no significant ABC transporter expression as tested by specific inhibitors), and these data were used for the correction of the data obtained in ABC transporter expressing cells. IC_50_ values were calculated by nonlinear regression analysis using the Origin2019 software (version 9.6.5.169, OriginLab Corporation, Northampton, MA, USA).

In addition to studies on the wild-type ABCG2 transporter, we have also examined the effects of drugs on the function of a relatively frequent (present in up to 20-35% of the population) polymorphic variant, the Q141K-ABCG2 (SNP rs2231142) transporter. In these experiments we measured Hoechst dye extrusion in HeLa cells stably expressing eGFP and the ABCG2 variants driven by the same promoter, the transgenic cells generated by the Sleeping Beauty transposon system. These cells were sorted for similar levels of eGFP, and ABCG2 expression was examined by a monoclonal (5D3) antibody-based flow cytometry assay (Q141K-ABCG2 cell surface expression was approximately 80% of the wild type ABCG2 expression. For details see Zámbó *et al.*[45]).

For the ABCG2 transporter inhibition assay, trypsinized HeLa cells expressing the ABCG2 WT or Q141K variant were incubated in a shaker for 20 minutes at 37°C in HPMI buffer (20mM HEPES, 132 mM NaCl, 3.5 mM KCl, 0.5 mM MgCl_2_, 5mM glucose, 1 mM CaCl_2_ [pH 7.4]) with 0.2 μM of Hoechst 33342 (H1399, Thermo Fisher Scientific, Waltham, MA, US) dye, in the presence of the tested drugs, or 1 μM of the specific ABCG2 inhibitor, Ko143 (3241, Tocris Bioscience, UK). Following incubation, the cells were put on ice and Hoechst dye fluorescence was measured using the violet laser (405 nm) and VL1 detector on the Attune Nxt Cytometer. Cells showing similar eGFP fluorescence were gated in the case of the WT and Q141K variants. The maximum inhibition was defined for each variant as the fluorescence difference of the samples incubated with and without Ko143, respectively. All tested drugs (ivermectin, remdesivir, ritonavir, lopinavir and Ko143) were dissolved in DMSO, and the same DMSO concentration (below 0.5%) was applied for all drug concentrations examined. Data analysis was performed using the Attune Nxt Cytometer Software v3.1.2. Two samples per condition and three biological replicates were examined in each experiment.

*For studying vesicular ATP-dependent transporter activity* of the ABC transporters, we used ABCB1/Pgp or ABCG2/BCRP containing HEK-293 cell membrane vesicles, as well as ABCC1/MRP1 containing Sf9 membrane vesicles, prepared by Solvo Biotechnology. Membrane vesicles (12.5 μg protein/sample) were incubated with transporter-specific substrates, for ABCG2 5 μM lucifer yellow (LY) as a fluorescent substrate, and the radiolabeled substrate probes, for ABCB1 1 μM N-methyl quinidine (NMQ), and for ABCC1 0.2 μM estradiol-glucuronide (ETGB), at 37°C for 10 min (ABCG2), 1 min (ABCB1) or 3 min (ABCC1), with or without 4 mM Mg-ATP, in 50 μL final volume. The known reference inhibitors of the transporters (1 μM Ko143 for ABCG2, 1 μM Valspodar for ABCB1, and 200 μM Benzbromarone for ABCC1) served as controls. Each test compound was dissolved in DMSO and 1 μL volume was added to the samples. After incubation, the samples were rapidly filtered and washed on filter plate (MSFBN6B10, Millipore, Burlington, MA, US). Accumulated substrates in the vesicles were dissolved by 100 μL of 10% SDS and then centrifuged into another plate. In case of fluorescent substrates 100 μL of fluorescence stabilizer was added to the samples (DMSO for LY). Fluorescence was measured in plate readers (Victor X3 and Enspire Perkin-Elmer, Waltham, MA, US) at appropriate wavelengths, while the activity of the radiolabeled substrate was measured by liquid scintillation counting (Perkin Elmer MicroBeta2 liquid scintillation counter, Perkin Elmer, Waltham MA). ABC transporter-related vesicular transport was calculated by subtracting passive uptake measured without Mg-ATP (or in the presence of Mg-AMP) from the values measured in the presence of Mg-ATP. No significant quenching effect of the test compounds was observed.

*ABC transporter ATPase activity* was measured in Sf9 membrane vesicles containing the respective human ABC transporters[46–50]. Briefly, human ABCB1/Pgp, or ABCG2/BCRP were expressed in Sf9 insect cells by using baculovirus vectors. For ABCG2/BCRP, the cholesterol level in the vesicles was adjusted to the level of mammalian membranes to obtain full activity [49]. Membrane vesicles were stored at −80°C and total protein content of the preparations was measured by Lowry method. For ABCB1/Pgp 20 μg/150 μL, and for ABCG2/BCRP 10 μg/150 μL vesicles were incubated with 3 mM Mg-ATP for 25 min at 37°C. The test drugs were dissolved in DMSO and added in 1 μL to the samples. ABC transporter-related ATPase activity was determined as vanadate sensitive ATPase activity, and reference activators (50 μM verapamil for ABCB1, and 5 μM quercetin for ABCG2) served as controls.

### OATP transporter assays

For studying the function of the *multispecific uptake transporters, OATP1A2 and OATP2B1*, we have used A431 human tumor cells overexpressing these human proteins and mock transfected A431 cells as controls, generated as previously described[47,51]. The interaction between OATPs and the potential anti-COVID-19 agents was investigated in an indirect assay[47] employing two recently described fluorescent dye substrates, pyranine for OATP2B1, and sulforhodamine 101 for OATP1A2[51,52], as transported fluorescence indicators of OATP function.

Briefly, A431 cell overexpressing OATP1A2 or OATP2B1[47] were seeded on 96-well plates in a density of 8×10^4^ cells /well in 200 μL DMEM one day prior to the transport measurements. Next day, the medium was removed and the cells were washed three times with 200 μL phosphate buffered saline (PBS, pH 7.4) and pre-incubated with 50 μL uptake buffer (125 mM NaCl, 4.8 mM KCl, 1.2 mM CaCl_2_, 1.2 mM KH_2_PO_4_, 12 mM MgSO_4_, 25 mM MES (2-(Nmorpholino) ethanesulfonic acid, and 5.6 mM glucose, pH 5.5 for OATP2B1 and pH 7.0 for OATP1A2) at 37°C with or without increasing concentrations of the tested compounds. Each test compound was dissolved in DMSO (that did not exceed 0.5% in samples), solvent controls were also applied. The reaction was started by the addition of 50 μL uptake buffer containing a final concentration of 20 μM pyranine (OATP2B1), or 0.5 μM sulforhodamine 101 (OATP1A2).

Thereafter, cells were incubated at 37°C for 15 min (OATP2B1) or 10 min (OATP1A2). The reactions were stopped by removing the supernatants, then the cells were washed three times with ice-cold PBS. Fluorescence (in 200 μL PBS/well) was determined employing an Enspire plate reader (Perkin Elmer, Waltham, MA) ex/em: 403/5 and 517/5 nm (pyranine) or 586/5 and 605/5 nm (sulforhodamine 101). OATP-dependent transport was calculated by extracting fluorescence measured in mock transfected cells. The experiments were repeated in three biological replicates. IC_50_ values were calculated by nonlinear regression analysis using GraphPad prism software (version 5.01, GraphPad, La Jolla, CA, US).

## 3. Results

### 3.1. Interaction of anti-COVID-19 drug candidates with ABCB1/MDR1/Pgp

In these experiments we have applied several independent *in vitro* methods to explore the drug interactions with this transporter. We have measured ABCB1-related fluorescent substrate (Calcein-AM) transport inhibition in intact model cells, inhibition of the vesicular transport of an established ABCB1 substrate N-methyl quinidine (NMQ) in mammalian cell membrane vesicles, and drug-dependent ABCB1-ATPase activity in isolated insect (Sf9) cell membranes (see Figure 1). In all cases previously well characterized cells or membranes with highly expressed ABCB1 levels were applied, thus the data can be compared to numerous similar studies in the relevant scientific literature.

**Figure 1.**
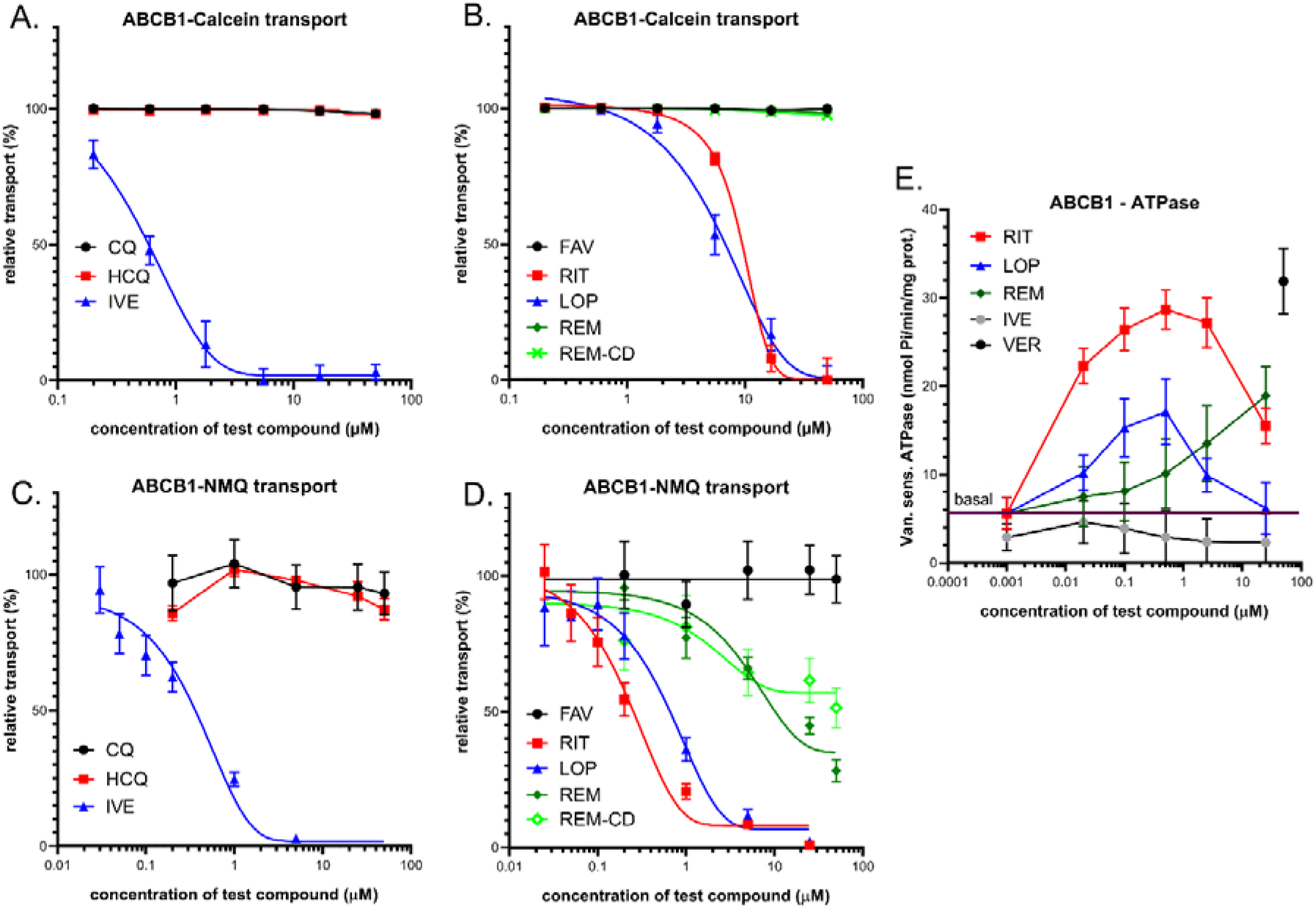
Panels A and B: Inhibition of ABCB1-mediated CaAM extrusion in intact ABCB1 expressing PLB-985 cells. Panel A: effects of ivermectin (IVE), chloroquine (CQ) and hydroxychloroquine (HCQ). Panel B: effects of lopinavir (LOP), ritonavir (RIT), favipiravir (FAV), remdesivir (REM) and cyclodextrin formulated remdesivir (REM-CD). Panels C and D: Inhibition of ABCB1-mediated N-methyl quinidine (NMQ) transport in the vesicular transport assay. Panel A: effects of ivermectin, chloroquine and hydroxychloroquine. Panel B: effects of lopinavir, ritonavir, favipiravir, remdesivir and cyclodextrin formulated remdesivir (REM-CD). Panel E: ABCB1-ATPase activity in isolated Sf9 membrane vesicles. Effects of ivermectin, lopinavir, ritonavir, and remdesivir. As a reference substrate, verapamil (VER) was used. The basal line represents ATPase activity level without the addition of any drug. Data on the graphs show the average of at least three independent experiments, +/− SD or SEM (panel E) values.

#### 3.1.1. A Transport assays in intact human PLB-985/ABCB1 cells1

ABCB1 is a high affinity active efflux transporter of a viability dye, Calcein-AM (CaAM). The cellular uptake of the non-fluorescent CaAM is strongly inhibited by the extrusion of this compound by the ABCB1 transporter, thus the cellular fluorescence of the intracellularly formed free Calcein is strongly reduced in cells expressing ABCB1[53]. Here we applied this widely used cellular assay (see https://www.solvobiotech.com/services/categories/dye-efflux-assays) to characterize drug interactions of the applied compounds in intact PLB-985 cells, expressing high levels of the ABCB1 protein. Maximum inhibition of the transporter was achieved by 0.25 μM tariquidar (TQ), a third-generation specific inhibitor of ABCB1. ABCB1 inhibition by the test compounds was estimated by their relative (%) inhibition compared to full inhibition by TQ.

As shown in Figure 1A and 1B, ivermectin caused a strong inhibition of the ABCB1-dependent CaAM extrusion at low micromolar concentrations (estimated IC_50_ 0.6 μM), while chloroquine and hydroxychloroquine had practically no effect on this ABCB1-dependent transport. Figure 1B documents the effects of lopinavir, ritonavir, favipiravir and remdesivir on the CaAM extrusion by ABCB1. Favipiravir, remdesivir and its cyclodextrin complex did not significantly inhibit the transport activity of ABCB1. In contrast, lopinavir and ritonavir caused a strong inhibition of the ABCB1-dependent CaAM extrusion at low micromolar concentrations (estimated IC_50_ values 6.3 and 8.4 μM, respectively).

#### 3.1.2. Vesicular transport studies in HEK/ABCB1 membrane vesicles

In these experiments we have studied the effects of the potential anti-COVID-19 compounds in a vesicular transport assay, by using inverted membrane vesicles prepared from HEK-293 cells expressing high levels of the ABCB1 transporter. In the inverted membrane vesicle measurements, labeled substrates and the investigated compounds are both applied on the cytoplasmic side of the membrane, that excludes most of the complex intracellular drug interactions. The modulation of the ATP-dependent vesicular uptake of a specific ABCB1 probe substrate directly reveals drug interactions with the transporter. In the present work we measured the uptake of 3H-N-methyl-quinidine (NMQ), a labeled low permeability amphipathic substrate of the human ABCB1/MDR1/Pgp[54,55]. Transport inhibition (%) of this labeled substrate by the test compounds is calculated by setting probe substrate transport in the absence of test compounds as 100%. As shown in Figure 1C, in ABCB1 expressing membrane vesicles ivermectin caused a strong inhibition of the ATP-dependent NMQ uptake at low micromolar concentrations (estimated IC_50_0.3 μM), while chloroquine and hydroxychloroquine had practically no effect on this transport activity. Figure 1D shows the effects of lopinavir, ritonavir, favipiravir and remdesivir on the transport activity of ABCB1. As documented, lopinavir and ritonavir are strong inhibitors of this transport (estimated IC_50_ values are 0.6 and 0.3 μM, respectively), remdesivir and remdesivir-SBECD are weak inhibitors of ABCB1 (IC_50_ above 20 μM), and favipiravir had no inhibitory effect.

#### 3.1.3. ABCB1-ATPase activity measurements in Sf9 membranes

Drug-stimulated ATPase activity of the ABCB1transporter is a well-documented functional assay to estimate the substrate or inhibitory features of various drugs. An ATP-dependent substrate transport is reflected in the activation of this specific ATPase activity in most cases, and in many cases drug-stimulated ABCB1-ATPase has been shown to correlate with the substrate affinity. In this assay most transported substrates show a biphasic curve: low concentrations stimulate, while higher concentrations inhibit the ATPase activity[46].

In these experiments we have measured the vanadate-sensitive (ABC transporter related) ABCB1-ATPase activity in membrane vesicles isolated from ABCB1 overexpressing Sf9 cells[46,49]. Sf9 cells have low intrinsic membrane ATPase activity, thus both the baseline and the drug-stimulated ATPase activity of the ABCB1 protein can be measured. High level stimulation of this ATPase activity can be achieved by 50 μM of verapamil, a transported substrate of ABCB1, serving as a positive control in this assay.

As shown in Figure 1, panel E, several tested compounds significantly increased the ABCB1-ATPase activity in this assay. Low micromolar concentrations of ritonavir (EC_50_ less than 0.1 μM) stimulated the ATPase activity up to the level of verapamil activation, and both lopinavir (EC_50_ about 0.05 μM) and remdesivir (EC_50_ about 10 μM) showed significant ABCB1-ATPase stimulation. Ritonavir and lopinavir showed maximal stimulation at around 0.5 μM, whereas remdesivir increased the ATPase activity only at 10-50 μM. At higher concentrations ritonavir and lopinavir showed inhibitory effects. Ivermectin showed only ABCB1-ATPase inhibition.

### 3.2. Interaction of anti-COVID-19 drug candidates with ABCC1/MRP1

#### 3.2.1. Transport assay in intact human cells – HL60/ABCC1 cells

ABCC1, similarly to ABCB1, is a high affinity active efflux transporter for Calcein-AM and the cellular fluorescence of free Calcein is also reduced in cells expressing ABCC1[56]. Therefore, we applied this assay to characterize the test drug interactions in intact HL60 cells expressing high levels of the ABCC1 protein. Maximum inhibition of the transporter was achieved by 10 μM of indomethacin (IM), a strong inhibitor of ABCC1. ABCC1 inhibition by the test compounds was estimated by their relative (%) inhibition compared to full inhibition by IM.

As shown in Figure 2A, ivermectin caused a strong inhibition of the ABCC1-dependent CaAM extrusion at low micromolar concentrations (estimated IC50 3.3 μM), while chloroquine and hydroxychloroquine had practically no effect. As documented in Figure 2B, lopinavir and ritonavir had strong inhibitory effects with estimated IC50 values of 10.7 μM and 7.7 μM, respectively. Favipiravir, remdesivir or its cyclodextrin complex did not significantly inhibit the cellular transport activity of ABCC1 (a slight inhibition by higher concentrations of remdesivir was observed).

**Figure 2.**
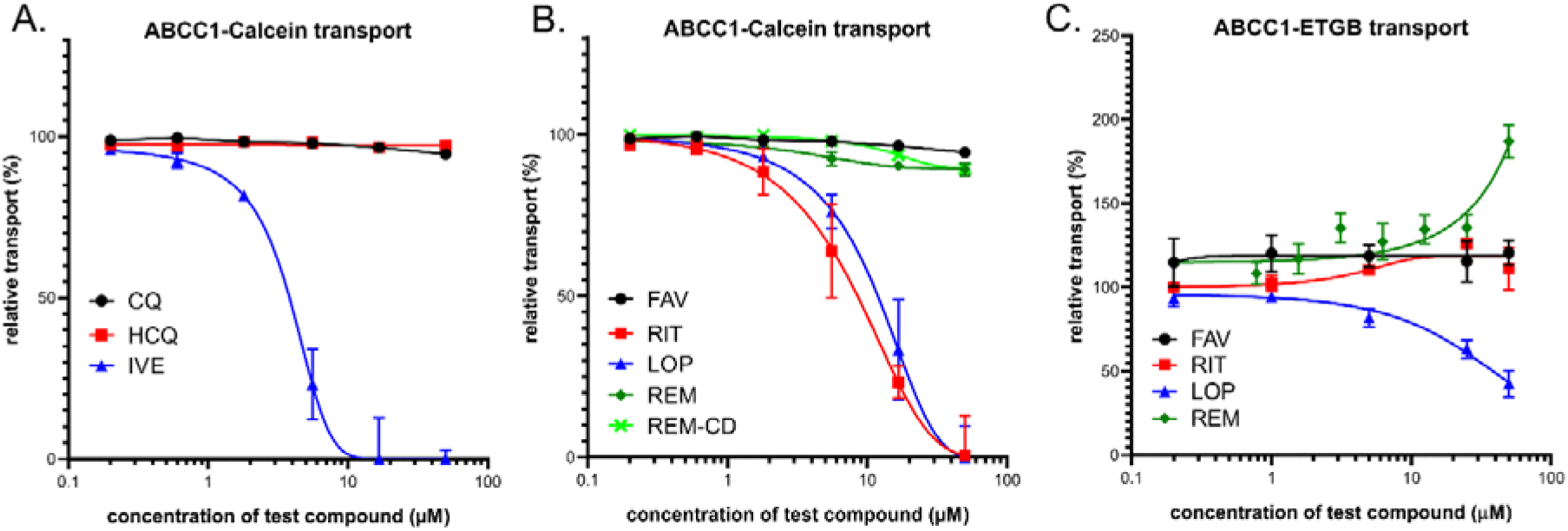
Panels A and B: Inhibition of ABCC1-mediated CaAM extrusion in intact HL60 cells. Panel A: effects of ivermectin (IVE), chloroquine (CQ) and hydroxychloroquine (HCQ). Panel B: effects of lopinavir (LOP), ritonavir (RIT), favipiravir (FAV), remdesivir (REM) and SEBCD-remdesivir (REM-CD).

#### 3.2.2. Vesicular transport studies in Sf9/ABCC1 membrane vesicles

Here we measured the effects of the test compounds in a vesicular transport assay, by using inverted membrane vesicles prepared from Sf9 cells expressing high levels of the ABCC1 transporter. We measured the uptake of ^3^H-estradiol-17b-glucuronide (ETGB), a labeled substrate of the human ABCC1/MRP1 transporter[57]. Relative inhibition (%) was calculated by setting probe substrate transport in the absence of test compounds as 100%.

As shown in Figure 2C, in ABCC1 expressing membrane vesicles lopinavir caused an inhibition of the ATP-dependent ETGB uptake in a dose-dependent manner with a maximum inhibition of 60 % at the highest applied concentration. Ivermectin inhibited the ABCC1-mediated ETBG accumulation with a maximum inhibition of 55 % at 25 μM concentration (not shown). Ritonavir, favipiravir, chloroquine and hydroxychloroquine had practically no effect on this transport activity (CQ and HCQ not shown), while, interestingly, remdesivir significantly stimulated the vesicular ETGB transport activity of ABCC1, although only at above 25 μM.

### 3.3. Interaction of anti-COVID-19 drug candidates with ABCG2

#### 3.3.1. Transport measurements in intact human cells – PLB/ABCG2 and HeLa/ABCG2

The ABCG2 protein does not transport Calcein AM, thus this assay cannot be used for assaying ABCG2 activity in intact cells. In contrast, several fluorescent dyes, including Hoechst 33342 (Hst), DyeCycle violet (DCV) or PhenGreen-SK diacetate (PG-DA) are actively extruded by the ABCG2 transporter[44,58–63], providing an opportunity for fluorescence-based cellular assays. Here we used both PhenGreen-SK diacetate and Hst dye for these measurements.

The PhenGreen (PG) assay is a recently patented method for an efficient ABCG2 transport measurement, as the non-fluorescent PhenGreen diacetate (PG-DA) is actively extruded by ABCG2 and the cellular fluorescence is correlated with ABCG2 transport activity[44]. Therefore, we applied this assay to characterize the test drug interactions in intact PLB-985 cells expressing high levels of the ABCG2 protein. Maximum inhibition of the transporter was achieved by 2.5 μM of Ko143, a high-affinity specific inhibitor of ABCG2. ABCG2 inhibition by the test compounds was estimated by their relative (%) inhibition compared to full inhibition by Ko143.

As shown in Figure 3A and B, ivermectin caused a strong inhibition of the ABCG2-dependent PG-DA extrusion at micromolar concentrations (estimated IC_50_ 3.1 μM). Lopinavir and ritonavir also had strong inhibitory effects, with estimated IC_50_ values of 13.1 μM and 8.3 μM, respectively. Favipiravir, chloroquine, and hydroxychloroquine had practically no effect on this ABCG2-dependent transport. Remdesivir and its complex are relatively weak inhibitors of the ABCG2-dependent PG-DA extrusion at micromolar concentrations (estimated IC_50_ values are greater than 40-50 μM.)

**Figure 3.**
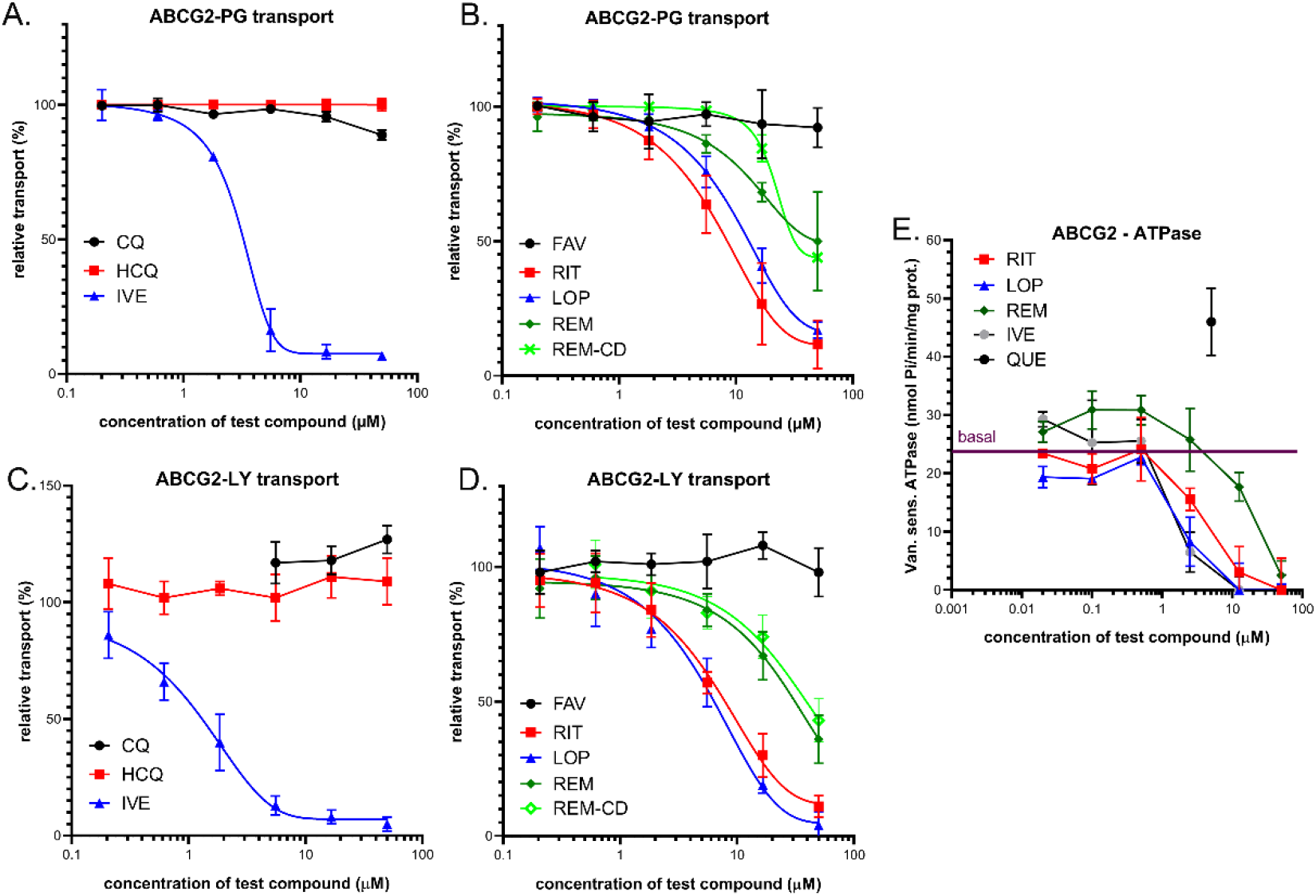
Panels A and B: Inhibition of ABCG2-mediated PG-AM extrusion in intact PLB-985 cells. Panel A: effects of ivermectin (IVE), chloroquine (CQ) and hydroxychloroquine (HCQ). Panel B: effects of lopinavir (LOP), ritonavir (RIT), favipiravir(FAV), remdesivir (REM) and SEBCD-remdesivir (REM-CD). Panels C and D: Inhibition of ABCG2-mediated lucifer yellow (LY) transport in the vesicular transport assay. Panel A: effects of ivermectin, chloroquine and hydroxychloroquine. Panel B: effects of lopinavir, ritonavir, favipiravir, remdesivir and SEBCD-remdesivir. Panel E: ABCG2-ATPase activity in isolated Sf9 membrane vesicles. Effects of ivermectin, lopinavir, ritonavir, and remdesivir. Maximum ATPase stimulation was obtained by 5 μM quercetin. Data on the graphs show the average of at least three independent experiments, +/− SD (Panels A and B) or SEM (panels C, D and E) values.

#### 3.3.2. Vesicular transport studies in HEK/ABCG2 membrane vesicles

The effects of the test compounds on ABCG2 activity were also measured in a vesicular transport assay, by using inverted membrane vesicles prepared from HEK-293 cells expressing high levels of the ABCG2 transporter. The ATP-dependent uptake of lucifer yellow (LY), a substrate of the human ABCG2/BCRP transporter[64] was measured, and inhibition of this fluorescent substrate uptake by the test compounds was compared to a full inhibition achieved by 1 μM of Ko143.

As shown in Figure 3, Panels C and D, in ABCG2 containing HEK-293 cell membrane vesicles, ivermectin caused a strong inhibition of the ATP-dependent LY uptake at low micromolar concentrations (estimated IC_50_ 1.1 μM), while chloroquine and hydroxychloroquine had practically no effect. Lopinavir, ritonavir, and remdesivir as well as remdesivir-cyclodextrin inhibited the transport activity of ABCG2 with estimated IC_50_ values of 4.2 μM, 7.5 μM and more than 20 μM, respectively.

#### 3.3.3. ABCG2-ATPase activity measurements in Sf9 membranes

Drug-stimulated ATPase activity of the ABCG2 transporter has also been measured in the present study. This functional assay may be used to estimate the substrate / inhibitory features of various drugs (see 1.C, and [50]). Here we measured the vanadate-sensitive ABCG2-ATPase activity in membrane vesicles isolated from ABCG2 overexpressing Sf9 cells[50], and a maximum stimulation of this ABCG2-ATPase activity was achieved by the transported substrate 5 μM quercetin, serving as a positive control in this assay. As shown in Figure 3, Panel E, ivermectin, ritonavir, lopinavir at low micromolar concentrations and remdesivir at higher concentrations (around 20-50 micromoles) significantly inhibited the baseline ABCG2-ATPase activity. None of the tested compounds showed significant ABCG2-ATPase stimulation effects, thus the assay was not informative regarding their potential transported substrate nature.

#### 3.3.4. Effect of the Q141K-ABCG2 polymorphism on the inhibitory potential of the test drugs

The very frequent (present in 12-35% in various populations) Q141K-ABCG2 polymorphic variant has been reported to have lower membrane expression levels and reduced transport activity in various assay systems[65]. Since ivermectin, lopinavir, ritonavir and remdesivir exerted well measurable inhibition on the ABCG2-dependent PG dye extrusion activity (see above), we have performed similar studies in HeLa cells stably expressing either the wild-type ABCG2 or the Q141K-ABCG2 variant[45]. In this case we measured the extrusion of the Hoechst 33342 dye, another transported substrate of ABCG2, and used again the specific inhibitor Ko143 to achieve full inhibition of the transporter.

As shown in Figure 4, ivermectin (A), ritonavir (B) and lopinavir (C) inhibited Hst dye extrusion by ABCG2 also in the HeLa cells, and remdesivir (D) had a slight effect at relatively high concentrations. An important finding was that Hst dye extrusion in HeLa cells expressing the Q141K-ABCG2 variant showed higher sensitivity to all drug inhibition.

**Figure 4:**
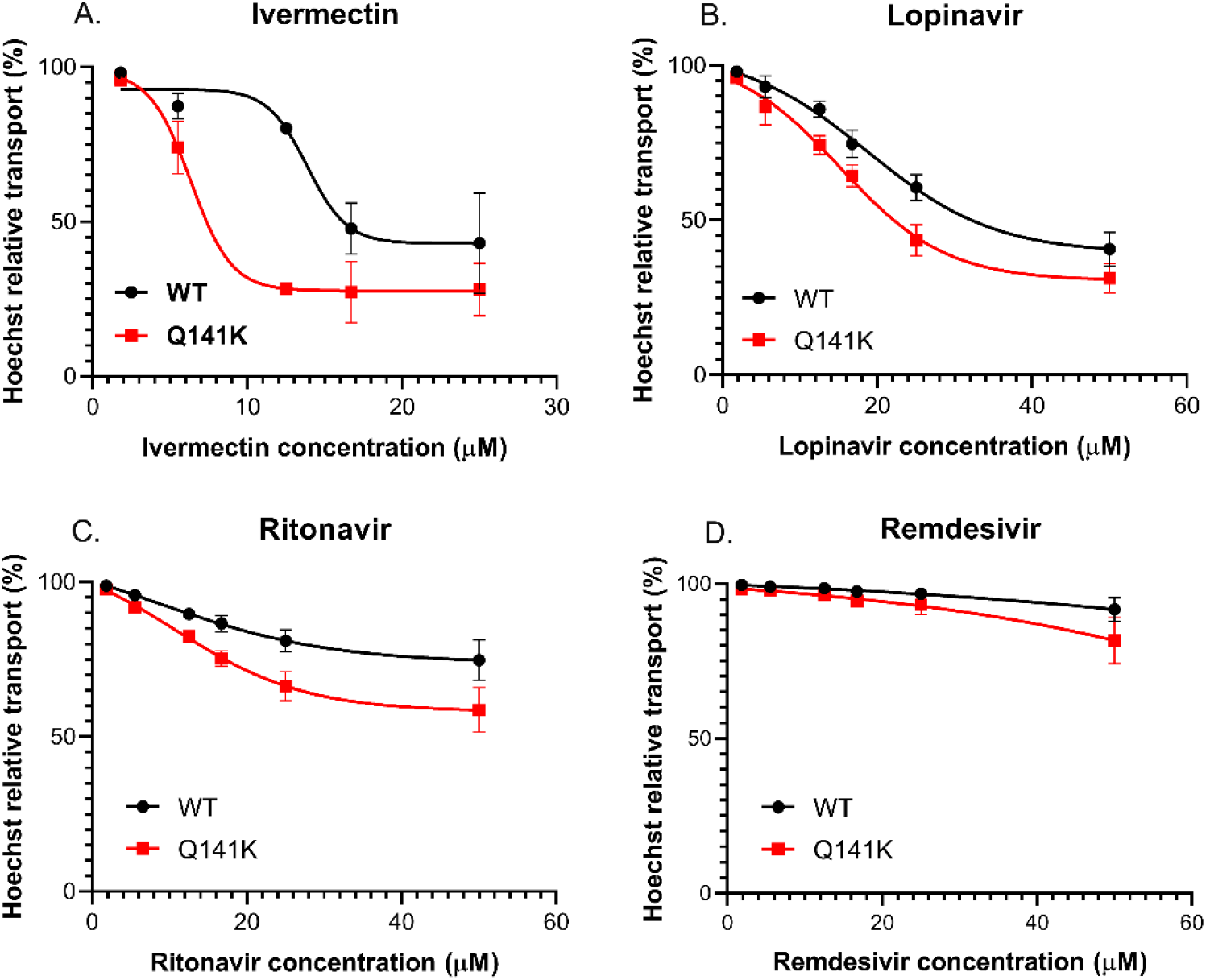
Hoechst 33342 dye extrusion in HeLa cells stably expressing the WT and Q141K variants of ABCG2. Different concentrations ivermectin (A), lopinavir (B), ritonavir (C) and remdesivir (D) were used to determine their effect on ABCG2 function. Data on the graphs show the average of three independent experiments, +/− SD values.

### 3.4. Interaction of anti-COVID-19 candidates with OATP1A2 and OATP2B1 transporters

In order to explore potential interaction between the potential anti-COVID-19 drugs and OATP1A2 or OATP2B1, the compounds were investigated in a fluorescence-based cellular transport assay recently developed by our laboratory[47,51]. Uptake of pyranine or sulforhodamine 101 as test substrates was measured in A431-OATP1A2 or A431-OATP2B1 cells, and mock transfected A431 were used as negative control. As shown on Figure 5, with the exception of favipiravir, all the compounds examined inhibited OATP1A2 function. Based on IC_50_ values, antivirals lopinavir, ritonavir, remdesivir, and the anti-parasitic ivermectin showed similar affinities towards OATP1A2. On the other hand, although still effective, chloroquine or hydroxychloroquine were 3-10-fold lower affinity inhibitors (see Table 2). Interestingly, cyclodextrin resulted in slightly decreased inhibitory potential of remdesivir on OATP1A2 function (Figure 5, and Table 2).

**Table 1.** Mechanism of action of the potential anti-COVID-19 drugs examined in this study

| Potential anti-COVID-19 compound | Mechanism of action |
| --- | --- |
| chloroquine | Antimalarial - endosomal pH increase |
| hydroxychloroquine | Antimalarial - endosomal pH increase |
| ivermectin | Antiparasitic - glutamate-gated chloride channel and a GABA receptor inhibitor |
| lopinavir | (HIV) protease inhibitor |
| ritonavir | (HIV) protease inhibitor |
| remdesivir | Viral RNA-polymerase inhibitor |
| favipiravir | Viral RNA-polymerase inhibitor |

**Table 2.** Summary of the transporter inhibition properties of the drugs examined. Approximate IC_50_ (μM) values were determined by nonlinear regression analysis of data shown in the Results section, using GraphPad prism software (version 5.01, GraphPad, La Jolla, CA, USA).

|  | Estimated transporter inhibition - IC50 (µM) |  |  |  |  |  |  |  |
| --- | --- | --- | --- | --- | --- | --- | --- | --- |
|  | ABCB1 |  | ABCC1 |  | ABCG2 |  | OATP - | cellular assays |
| Potential anti-COV-19 compounds | cellular assay | vesicular assay | cellular assay | vesicular assay | cellular assay | vesicular assay | OATP1A2 | OATP2B1 |
| chloroquine | - | - | - | - | - | - | 17.0 | 119 |
| hydroxychloroquine | - | - | - | - | - | - | 18.9 | 84 |
| ivermectin | 0.6 | 0.3 | 3.3 | 25 | 3.1 | 1.1 | 5.2 | 8.6 |
| lopinavir | 6.3 | 0.6 | 10.7 | 10 | 13.1 | 4.2 | 1.5 | 1.0 |
| ritonavir | 8.4 | 0.3 | 7.7 | - | 8.3 | 7.5 | 2.3 | 1.4 |
| remdesivir | - | >20 | - | - * | >50 | >50 | 3.8 | 3.8 |
| remdesivir-SBECD | - | >20 | - | NA | >50 | >50 | 6.1 | 5.6 |
| favipiravir | - | - | - | - | - | - | - | - |

\* Stimulation of substrate transport

**Figure 5.**
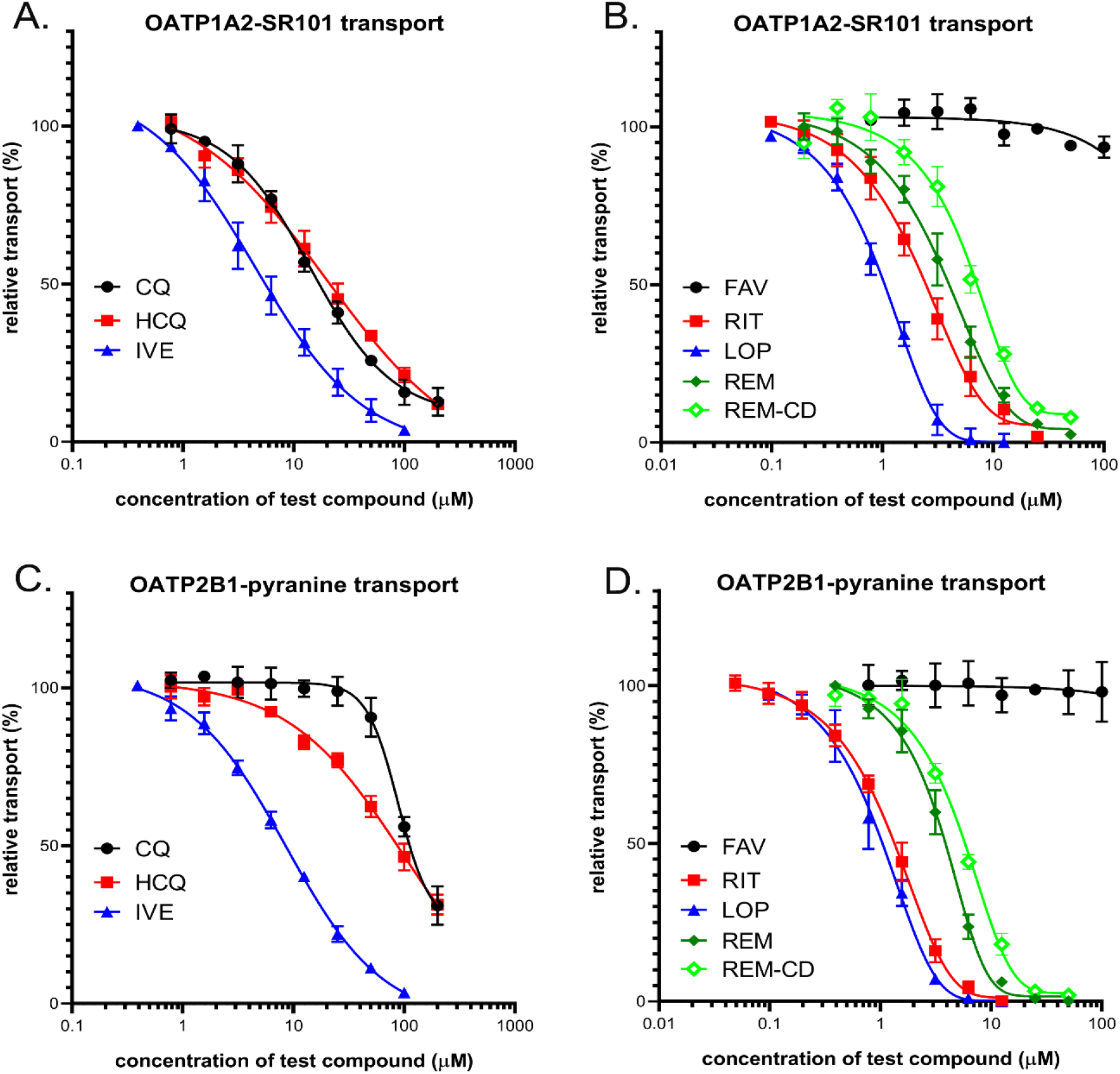
Panels A and B: Inhibition of OATP1A2-mediated sulforhodamine101 (SR101) uptake by potential antiviral compounds. Uptake of 0.5 μM SR101 was measured in A431-OATP1A2 cells seeded on 96-well plates for 10 minutes in the presence of increasing concentrations of ivermectin (IVE), chloroquine (CQ), hydroxychloroquine (HCQ), favipiravir(FAV), lopinavir (LOP), ritonavir (RIT), remdesivir (REM) and SEBCD-remdesivir. Fluorescence was measured in an Enspire plate reader. Transport was determined by subtracting fluorescence in A431-mock cells. Uptake rates were expressed as percent of the uptake measured in the absence of any inhibitor. Data on the graphs show the average of at least three independent experiments, +/− SD values. Panels C and D: Inhibition of OATP2B1-mediated pyranine uptake by different antiviral compounds. Uptake of 20 μM pyranine was measured in A431-OATP2B1 cells seeded on 96-well plates for 15 minutes in the presence of increasing concentrations of the tested compounds. Uptake rates were determined as in Fig 5A. Averages obtained from at least three biological replicates. +/− SD values are shown.

In the case of OATP2B1, similar inhibitory potency was observed for the antiviral compounds as in the case of OATP1A2. Lopinavir and ritonavir were the highest affinity inhibitors with IC_50_ values of 1.0 and 1.4 μM, respectively, and remdesivir, remdesivir-cyclodextrin complex and ivermectin had lower inhibitory effects with IC_50_ values of 3.8 μM, 5.6 μM, and 8.6 μM, respectively. Chloroquine and hydroxychloroquine exerted only modest inhibition of OATP2B1 activity, and favipiravir showed no interaction with this transporter.

## 4. Discussion

Multispecific drug and xenobiotic transporters play a major role in the pharmacokinetics of numerous pharmacological agents, and any new drug candidates have to be tested for interactions with the key transporters in this regard[66,67]. However, several drugs rapidly repurposed in the past months for potential anti-COVID-19 activity, may not have been analyzed in detail for these interactions. This lack of information is making their clinical use, especially in combination with other pharmacological agents, potentially dangerous for the patients. Since the SARS-CoV-2 virus infection causes severe clinical symptoms especially in elderly patients and/or in those with existing co-morbidities, the ADME-Tox properties and the potentially harmful drug-drug interactions of the repurposed drugs may have major relevance in these cases. In addition, there are no current methods to correctly estimate the transporter-drug interactions by any *in silico* methods, and only detailed *in vitro* studies may answer these questions. Therefore, in the present study we have examined the interactions of the repurposed drug candidates with the key multispecific drug transporters with the hope that our results help the clinical applications in COVID-19 treatment.

**ABC transporters** play a key role in the pharmacokinetics of numerous pharmacological agents, and they, especially ABCB1 and ABCG2 should be tested in early phases of drug development[66]. As shown in the results section and in the summary Table 2, regarding the three ABC multispecific transporters examined, we found that **chloroquine, hydroxychloroquine and favipiravir** showed practically no potentially relevant interaction with any of these transporters. In contrast, **ivermectin** was found to exert a strong inhibitory effect both in the case of the ABCB1, ABCC1, and ABCG2 transporter. Some of these interactions have already been explored and, especially ivermectin inhibition of the human and animal Mdr1/Pgp/ABCB1 has been studied in detail[8,9,68]. As we show here, in all kinds of assays a strong inhibition of both ABCC1 and ABCG2 transport activity was also observed by ivermectin, emphasizing the potentially dangerous effects of this compound at higher doses or at impaired transporter function.

The antiviral protease inhibitors **lopinavir and ritonavir** had a significant inhibitory effect on the three ABC transporters examined here, and a strong inhibition was observed for ABCB1/Pgp (Table 2). However, ABCC1 and ABCG2 were also inhibited by these compounds in potentially relevant concentrations. As suggested by the relevant FDA information (https://www.accessdata.fda.gov/drugsatfda_docs/label/2007/021226s018lbl.pdf, https://www.ema.europa.eu/en/documents/scientific-discussion/kaletra-epar-scientific-discussion_en.pdf), lopinavir in patients may reach plasma concentrations of about 13 μM, while ritonavir may peak at about 1 μM, therefore our in vitro data indicate that drug-drug interactions should be considered.

**Remdesivir** was found to be a relatively weak inhibitor of all the three ABC transporters, although the ABCB1/Pgp inhibition with an IC_50_ of about 20 μM, observed in the vesicular transport assay, may be relevant under certain treatment conditions (reported peak plasma concentrations of remdesivir about 5 μM). Interestingly, remdesivir significantly stimulated the vesicular transport of a test substrate ETGB by the ABCC1 transporter, thus an allosteric effect of remdesivir on this transporter should be considered.

The experiments performed in intact cells expressing the polymorphic ABCG2 transporter variant, Q141K, with impaired membrane localization and transport activity, suggest that at lower transporter functions, some of the weak inhibitors may have a clinically important effect. In this case, both lopinavir and ritonavir exerted a significantly stronger inhibition of this drug transporter variant. Considering that the allele encoding the ABCG2-Q141K variant (rs2231142) has an incidence of about 30% in the Asian population, this information is especially important for clinical interventions in these countries.

In our studies we have also examined the membrane ATPase activity of the ABCB1 and the ABCG2 transporters in isolated membranes. These assays may help to decipher the substrate vs. inhibitor nature of the test compounds. Although these results provide only a tentative answer in this regard, we found that the ABCB1-ATPase activity is significantly stimulated by low concentrations of lopinavir, ritonavir and remdesivir, indicating that these drugs may be transported substrates. We observed no such stimulation in the case of the ABCG2 transporter by these drugs.

In addition to the ABC multidrug transporters, the role of drug transporting OATPs in pharmacokinetics and drug-drug or food-drug interactions is increasingly recognized [66]. While OATPs, 1B1 and 1B3 are liver-specific proteins, OATP1A2 and OATP2B1 are highly expressed in the endothelial cells of the BBB, therefore these transporters are key modulators of the entry of their drug substrates into the central nervous system[41,69,70]. In addition, OATP2B1 also influences the absorption of its substrates from the intestine[40].

In the current study we have investigated the interactions between **OATP1A2 and OATP2B1** and drugs repurposed to treat COVID-19 disease. We found high affinity interactions between the antiviral protease inhibitors **lopinavir and ritonavir** with both OATP1A2 and OATP2B1. These interactions have already been documented[71–74], moreover, lopinavir and ritonavir are also inhibitors of OATP1B1 and OATP1B3, the key transporters in the liver[71]. The IC_50_ values obtained in our study are in harmony with those observed in Tupova *et al.*[72] and Kis *et al.*[73]. Moreover, a study[74] showed that lopinavir is transported by OATP1A2, thus OATP1A2 probably affects lopinavir absorption and blood to brain entry. **Ivermectin**, as reported earlier[75], also inhibited OATP2B1, although in our experiments with higher affinity than that found by Karlgren *et al.* (20 μM 39%-this may be explained by the different test substrates used). However, here we document the first time that ivermectin inhibits OATP1A2 function. Our experiments confirmed the relatively weak interactions between **chloroquine and hydroxychloroquine** with OATP1A2 and OATP2B1, and the observed IC_50_ values harmonize with that found by others[76,77]. We observed no interaction of **favipiravir**with the two OATPs examined here.

There are no data available for the interaction of **remdesivir**, currently a major anti-COVID-19 drug candidate, with OATP1A2 and OATP2B1. Based on our data, showing a high-affinity interaction of remdesivir with both transporters, and considering the plasma concentrations of this drug (peak values in 3-5 μM[78,79] remdesivir may significantly interfere with the pharmacokinetics of OATP1A2 and OATP2B1 drug substrates. Our current methods do not distinguish between transported and non-transported inhibitors, thus further investigations are needed to decipher whether OATP1A2 and OATP2B1 can mediate the uptake of remdesivir.

From the currently developed anti-COVID-19 drugs tested in our study, favipiravir is the most promising candidate to avoid unexpected drug-drug interactions. Favipiravir, similarly to that found for the multispecific ABC transporters located in the tissue barriers (see above), did not inhibit the function of the investigated OATPs, therefore this compound is not expected to influence OATP mediated drug pharmacokinetics.

As a summary, our current data, summarized in Table 2, provide a detailed *in vitro* quantitative analysis on the interaction of the anti-COVID-19 agents with the key human multispecific drug transporters. These *in vitro* assays, which should be followed by careful clinical pharmacokinetic studies, may provide a strong warning against the compassionate use of some of these agents, especially when other relevant drugs are also applied to the patient, or in cases of endogenously impaired transporter activity.

## Author Contributions

Á.T., Cs. Ö-L. and B. S. wrote the draft, Cs.A., O.M. E.Sz. É.B. Á.T., and Gy. V. performed the experiments and critically revised the manuscript. All authors read and approved the final version.

## Acknowledgments

The 5D3 antibody was a kind gift of B. Sorrentino, St. Jude Children Hospital, Memphis, USA, remdesivir and remdesivir-SBECD were kind gifts of Lajos Szente, Cyclolab Ltd, Budapest, Hungary.

